# Predicting Intra- and Intertypic Recombination in Enterovirus 71

**DOI:** 10.1101/445783

**Authors:** Andrew Woodman, Kuo-Ming Lee, Richard Janissen, Yu-Nong Gong, Nynke Dekker, Shin-Ru Shih, Craig E. Cameron

**Affiliations:** department of Biochemistry and Molecular Biology, The Pennsylvania State University, 201 Althouse laboratory, University Park, PA 16802.; Research Center for Emerging Viral Infections, Chang Gung University, Taiwan.; Dept. of Bionanoscience; Kavli Institute of Nanoscience; Delft; South-Holland; 2629HZ; The Netherlands.; Department of Medical Biotechnology and Laboratory Science, College of Medicine, Chang Gung University, Taoyuan, Taiwan.; Department of Laboratory Medicine, Linkou Chang Gung Memorial Hospital.; Research Center for Chinese Herbal Medicine, Research Center for Food and Cosmetic Safety, and Graduate Institute of Health Industry Technology, College of Human Ecology, Chang Gung University of Science and Technology, Taoyuan, Taiwan.

**Keywords:** EV-A71, Recombination, Replicative recombination, Copy-choice, Predictive, Conservation in mechanism(s)

## Abstract

Enteroviruses are well known for their ability to cause neurological damage and paralysis. The model enterovirus is poliovirus (PV), the causative agent of poliomyelitis, a condition characterized by acute flaccid paralysis. A related virus, enterovirus 71 (EV-A71), causes similar clinical outcomes in recurrent outbreaks throughout Asia. Retrospective phylogenetic analysis has shown that recombination between circulating strains of EV-A71 produces the outbreak-associated strains which exhibit increased virulence and/or transmissibility. While studies on the mechanism(s) of recombination in PV are ongoing in several laboratories, little is known about factors that influence recombination in EV-A71. We have developed a cell-based assay to study recombination of EV-A71 based upon previously reported assays for poliovirus recombination. Our results show that: (1) EV-A71 strain-type and RNA sequence diversity impacts recombination frequency in a predictable manner that mimics the observations found in nature; (2) recombination is primarily a replicative process mediated by the RNA-dependent RNA polymerase (RdRp); (3) a mutation shown to reduce recombination in PV (L420A) similarly reduces EV-A71 recombination suggesting conservation in mechanism(s); and (4) sequencing of intertypic recombinant genomes indicates that template-switching is by a mechanism that requires some sequence homology at the recombination junction and that the triggers for template-switching may be sequence independent. The development of this recombination assay will permit further investigation on the interplay between replication, recombination and disease.

## Introduction

The Enterovirus genus in the family *Picornaviridae* currently consists of 15 species. Outside of rhinoviruses, the enteroviruses responsible for human mortality and morbidity fall specifically into groups A, B, C and D (*1*). This group of viruses, typified by poliovirus, has a 7.5 kb positive-sense RNA genome that encodes a single polyprotein that is flanked by non-coding regions (NCR). The polyprotein is co- and post-translationally processed by virus-encoded proteases to generate the structural proteins (VP4, VP2, VP3 and VP1), which assemble to form the icosahedral capsid and the non-structural proteins (2A^pro^, 2B, 2C, 3A, 3B^VPg^, 3C^pro^ and 3D^pol^) that mediate replication of the virus genome *(2, 3).*

RNA viruses, like those found in the Enterovirus genus, exist as a viral quasispecies as a consequence of misincorporations by their error-prone RNA-dependent RNA polymerases (RdRp) during genome replication *(4,5)*. In addition, recombination enables exchange of genetic material through a ‘ copy-choice’ mechanism in which the viral RdRp along with the nascent RNA switches templates during replication creating hybrids between two viruses replicating in the same cell *(6-8).* As well as being a driver of genetic variation, it is proposed that recombination may have evolved in order to ‘ rescue’ genomes from deleterious mutations that accumulate during error-prone replication *(9, 10).*

Enterovirus 71 (EV-A71), a member of the species A group, is an important neurotropic enterovirus for which there is currently no effective therapy or vaccine, and manifests most frequently as a childhood illness known as hand, foot, and mouth disease (HFMD) (11). However, acute EV-A71 infection can also be associated with flaccid paralysis, myocarditis or even fatal encephalitis *(11, 12).* EV-A71 variants have been classified into three groups (GgA, GgB, and GgC) and recombination has been linked to the founding of each subgroup lineage *(13).* More importantly, co-circulation of the species A EV-A71 and Coxsackievirus A16 (CV-A16) viruses has been associated with large-scale outbreaks of HFMD *(14, 15).* Sequence analysis of clinical isolates obtained since 2008 from patients with fatal neurological symptoms has demonstrated that these cases are mainly due to subgenogroup C4 of EV-A71, which was previously identified as an EV-A71 / CV-A16 recombinant virus *(1, 16).* In related enteroviruses, recombinant forms (RF), defined by serotype according to their capsid proteins, have been shown to emerge, prevail, and then disappear in temporal epidemiological surveys of globally distributed serotypes *(13, 17, 18).* In many of these examples, the recombinants are pathogenic.

It is evident that recombination is a critical driver of virus evolution with medically important consequences. While the triggers and mechanisms of recombination in PV are starting to be understood *(10, 19-21)*, the ability to predict the likelihood of a recombination event between circulating viruses of public health relevance has not been available. We wanted to use the cell-based approaches that have been developed to study recombination in PV as a tool to test whether recombination events for EV-A71 in cell-culture mimic what is observed in nature. To be able to predict a recombination event between circulating strains of EV-A71 would clearly be beneficial not only to the scientific community but to populations as a whole. In order to study recombination in EV-A71 we have developed a robust, reproducible cell-based assay. The established *in vitro* assay based upon the previously reported assays for PV *(19-21)* consists of two genomes each containing a different deleterious (and non-reverting) modification that prevents the production of viable progeny. The initial donor template in our assay is an EV-A71 C2 strain sub-genomic replicon *(22)*, which is replication competent but does not encode capsid proteins (Fig. 1A). The second genome, or acceptor template, has had a region that encompasses the active-site of the RdRp removed (Δ3D) making it replication incompetent (Fig. 1A). We show that co-transfection of human embryonic rhabdomyosarcoma (RD) cells leads to a recombination event between the non-structural protein encoding regions of the sub-genomic replicon (the polymerase donor template) and the structural encoding region of the Δ3D EV-A71 genome (acceptor template). In addition, our results indicate that recombination in EV-A71 is primarily a replicative process and therefore different from previously described non-replicative recombination in the related poliovirus (PV) *(23, 24)*. Current investigations of PV recombination have shown that targeted mutations to the RdRp of the donor and acceptor templates can significantly alter the yield of recombinant virus *(20, 21)*. We introduced a similar conserved mutation (L420A in PV, L421A in EV-A71) into our cell-based recombination assay and show a significant fall in recombinant yield. The same mutation had no impact on replication but led to an EV-A71 virus population that was ultra-sensitive to the antiviral Ribavirin. Crucially, we tested the biological relevance of our assay by expanding it to include two additional circulating strains of EV-A71: a strain known to recombine (C4), and a strain that is not recombinogenic (B5). Our results show important significant differences in viable recombinant virus yield that mimic observations found in nature. Limited sequencing of recombinant genomes suggested that no sequence motif acted as a trigger for recombination, but it did show that RNA sequence complementarity at the recombination site may be important, adding support to the widely accepted ‘ copy-choice’ mechanism of template switching. We believe that this assay can be used as a tool to predict the likelihood of recombination between current circulating EV-A71 strains.

**Figure 1.**
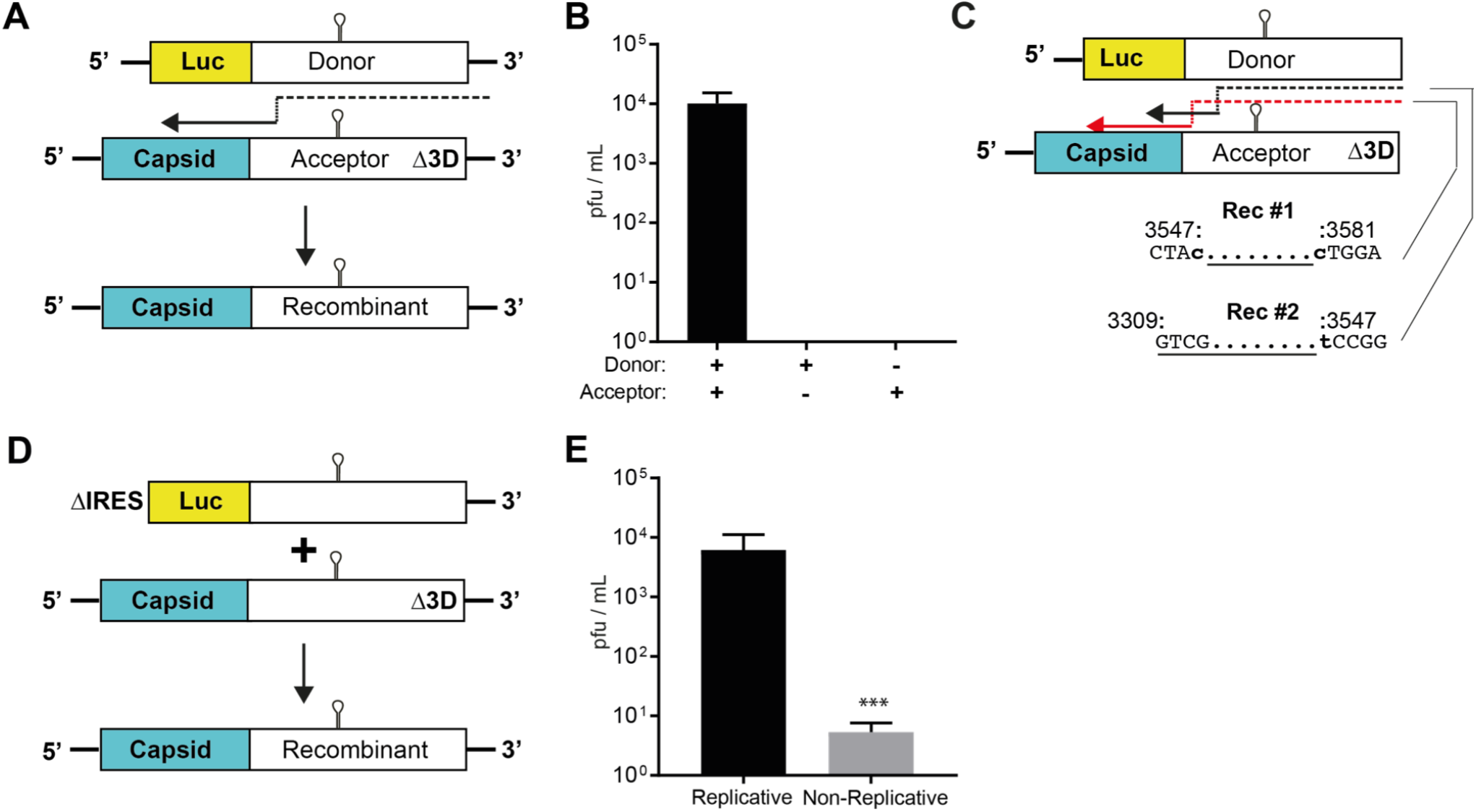
Enterovirus 71 (EV-A71) recombination in RD cells is primarily replicative. (**A**) Cell-based EV-A71 recombination assay. C2-strain firefly luciferase-encoding sub-genomic replicon (donor) and full length EV-A71 C2-MP4 strain genome (acceptor) carrying a lethal deletion of the 3D^po1^ region are co-transfected in equimolar ratio in RD cells. A fully functional virus genome can be produced via an RdRp template switch from donor to acceptor (indicated by dashed black arrow). (**B**) Only upon co-transfection, replication-competent virus can be generated (pfu/mL ±SD; *n = 3)* (**C**) Example sequences of plaque-purified recombinant virus from C2/C2 (left panel). Dashed arrows indicate predicted paths of viral RdRp upon template switching. Numbering refers to position upon the acceptor templates. Lower case, bold nucleotides indicate the 5’ and 3’ boundaries of recombination. Underlined sequences indicate region of homology. (**D**) Non-replicative recombination assay. IRES-deletion of the C2 donor template inhibits translation. Acceptor template remains the same as (A). Viable virus will only be produced via cell mediated event. **(E)** Yield of recombinant virus (pfu/mL ±SD; *n = 3)* originating from transfection in equimolar ratio of replicative and non-replicative partners. Statistical analyses were performed using unpaired, two-tailed t-test(***significance level p = 0.0001).

## Results

### Development of an EV-A71 cell-based recombination assay

In order to predict recombination events in EV-A71 an experimental system is required. The study of viral factors that modulate enterovirus recombination have benefitted from the recent development of recombination specific cell-based assays in PV that use parental templates that are only able to produce viable virus via recombination *(10, 19-21)*. A suitable ‘ donor’ template for the assay was the already established subgenogroup C2 EV-A71 (TW/2231/98) sub-genomic replicon where a Firefly luciferase reporter replaces the entire P1 region *(22)* (Fig. 1A). We modified the replicon by engineering a hammerhead ribozyme immediately 5’ of the internal ribosomal entry sequence (IRES), a change that would ensure an authentic genomic sequence following *in vitro* RNA transcription *(25)*. The modification led to a significant improvement in replication in RD cells with a 3log_10_ increase in reporter signal (a surrogate marker for genome replication) when compared to the unmodified replicon (Fig. S1). One recently used assay for the study of PV recombination, known as ‘ CRE-REP’ *(19)* uses an acceptor template that has characterized mutations within the 2C OriI stem-loop that inhibits positive-sense RNA synthesis *(26,27)*. The EV-A71 OriI has not been fully characterized, so similar mutations to the predicted stem-loop in 2C were not considered. In addition, mutations to the OriI of the related CVB3 virus have been shown to revert, and/or produce virus with 5’ RNA truncations *(28, 29)*. The ‘ acceptor’ template in our assay was the EV-A71 C2-MP4 strain *(30)* that had a region removed within the 3D-coding of the genome that encompasses the active site of the RdRp (EV-A71Δ3D), similar to an acceptor template used in a PV model for recombination *(21)*. Following co-transfection of donor and acceptor RNA templates into permissive cells, a viral-RdRp mediated template-switch from donor to acceptor may produce a fully-functional recombinant genome (Fig. 1A). Cell-based studies on the dynamics of EV-A71 replication are primarily carried out in RD or African green monkey (Vero) cells, which are susceptible to EV-A71 infection due to the expression of the receptor, SCARB2 *(31)*. As enterovirus recombination has been shown to be a replicative process *(10, 19-21)*, we wanted to select a cell-type for any EV-A71 recombination assay that would be optimal for replication of the donor replicon template. We initially quantified the luciferase signal, a marker for RNA replication, at 8 hours following transfection of RD and Vero cells with the EV-A71 C2 replicon RNA. Results indicated that Vero cells were sub-optimal for any future recombination assays as the luciferase signal was significantly lower (near 2Log_10_) than that produced in the RD cell line (Fig. S2). A subsequent co-transfection of RD cells with donor and acceptor RNA templates in equimolar ratios yielded viable recombinant virus which was quantified at 8.8 x 10^3^ pfu/mL (± 2 x 10^3^) (Fig. 1B), an amount in line with previous studies that used PV as a model *(19-21)*. Transfection of the donor and acceptor RNA templates alone produced no viable virus (Fig. 1B). To ensure that the observed virus was recombinant and to gain insight into the location of template switching, individual viruses were plaque purified and subjected to RT-PCR sequence analysis. Both donor and acceptor templates are derived from the EV-A71 subgenogroup C2 parental strains and share high sequence similarity (99.3% at the RNA level). Precise identification of the site of recombination would therefore be difficult. In two shown examples (Fig. 1C), the location of recombination was shown to fall within the 2A region. The first recombinant has a junction that falls within a 34-nucleotide window of shared homology between donor and acceptor templates. Similarly, the second recombinant has a junction that falls within a larger 248-nucleotide window. Importantly, all isolated sequences were recombinant, validating the experimental approach.

### EV-A71 recombination is primarily a replicative process

We generated an additional parental genome that would be unable to replicate in order to confirm whether the recombination we were observing was a replicative process and therefore different from non-replicative recombination *(32-35)*. We removed the entire IRES sequence from the EV-A71 C2 sub-genomic replicon producing the C2-ΔIRES-replicon template. This inhibits translation of the viral polypeptide, ensuring no active RdRp is produced (Fig. 1D). The second RNA partner in this ‘ non-replicative’ assay was the same acceptor template shown in figure 1A (EV-A71Δ3D). Importantly, both templates carried the relevant coding sequences that would produce a viable virus if a non-replicative mechanism of recombination occurred (Fig. 1D). In a side-by-side experiment with the replicative recombination assay, equimolar ratios of replicative partners and the newly developed non-replicative partners were transfected into RD cells. Quantification of virus 60 hours post transfection showed that the replicative partners were able to produce significantly more viable recombinant virus (5.4 x 10^3^ pfu/mL ± 2 x 10^3^) (Fig. 1E). In comparison, the non-replicative assay produced (~ 10 pfu/mL). This result is highly suggestive that recombination in EV-A71 is primarily a replicative process that is RdRp mediated, similar to that observed for PV *(10, 19-21)*.

### An RNA-dependent RNA polymerase (RdRp) mutation impairs recombination but not replication

Current investigations of PV recombination have shown that targeted mutations to the RdRp can significantly affect the yield of recombinant virus *(10, 19-21)*. A Lys-420-Ala change within the RdRp coding region has been shown to significantly inhibit recombination in a PV model (21). The Lys residue is conserved in the prototype strains of EV-A, B, C and D (position 421 of the RdRp coding region in EV-A71). We therefore reasoned that if the mechanism(s) of recombination were conserved between enterovirus species, a similar modification to the RdRp of the donor template might inhibit recombination in the newly developed EV-A71 recombination assay. We first introduced the L421A modification into the EV-A71 C2-MP4 full-length clone and quantified virus production following transfection of RNA into RD cells. No significant difference in virus yield was observed when compared to wild-type (Fig. 2A). In addition, luciferase signal ± guanidine hydrochloride of the L421A C2 replicon template was similar to wild-type replicon at 8 hours post-transfection (Fig. 2B). Taken together, the two experiments demonstrated that the L421A variant has no negative impact upon replication, an important pre-requisite for interpretation of any recombination assay. We then carried out a recombination assay to investigate the impact of the L421A mutation on virus yield. Following transfection of wild type parental RNA, a recombinant yield of 8.6 x 10^3^ pfu/mL (± 1 x 10^3^) was observed. In contrast, when the L421A mutation was present in the RdRp donor template the yield of recombinant was significantly reduced by more than 10-fold measuring an average of 5.3 x 10^2^ pfu/mL (± 2 x 10^2^) (Fig. 2C). Recombination has been proposed as an adaptive mechanism that generates combinations of beneficial mutations and/or removal of deleterious mutations that may appear in a population of viruses following replication *(36, 37)*. If this hypothesis is correct, then a virus population that is unable to ‘ purge’ deleterious mutations via recombination should be highly susceptible to mutagenic compounds such as nucleoside analogues. Indeed, a population of PV carrying the L420A mutation has been shown to be highly susceptible to the mutagen Ribavirin *(21)*. Kempf *et al*., have proposed that this is not related to RdRp fidelity but rather a direct result of inhibiting recombination. We tested an EV-A71 population carrying the similar L421A mutation. RD cells were infected at a MOI of 0.1 in the presence of increasing concentrations Ribavirin. Viable virus was then quantified by pfu and normalized to an untreated control. Results showed that concentrations of Ribavirin > 200 μM led to a significant decrease in the viability of the L421A EV-A71 population compared to wild-type (Fig. 2D). Importantly, the results with the L421A variant show that recombination in EV-A71 is replicative process and potentially indicate that the mechanism(s) of recombination and the consequences thereof in enteroviruses may indeed be conserved across species groups.

**Figure 2.**
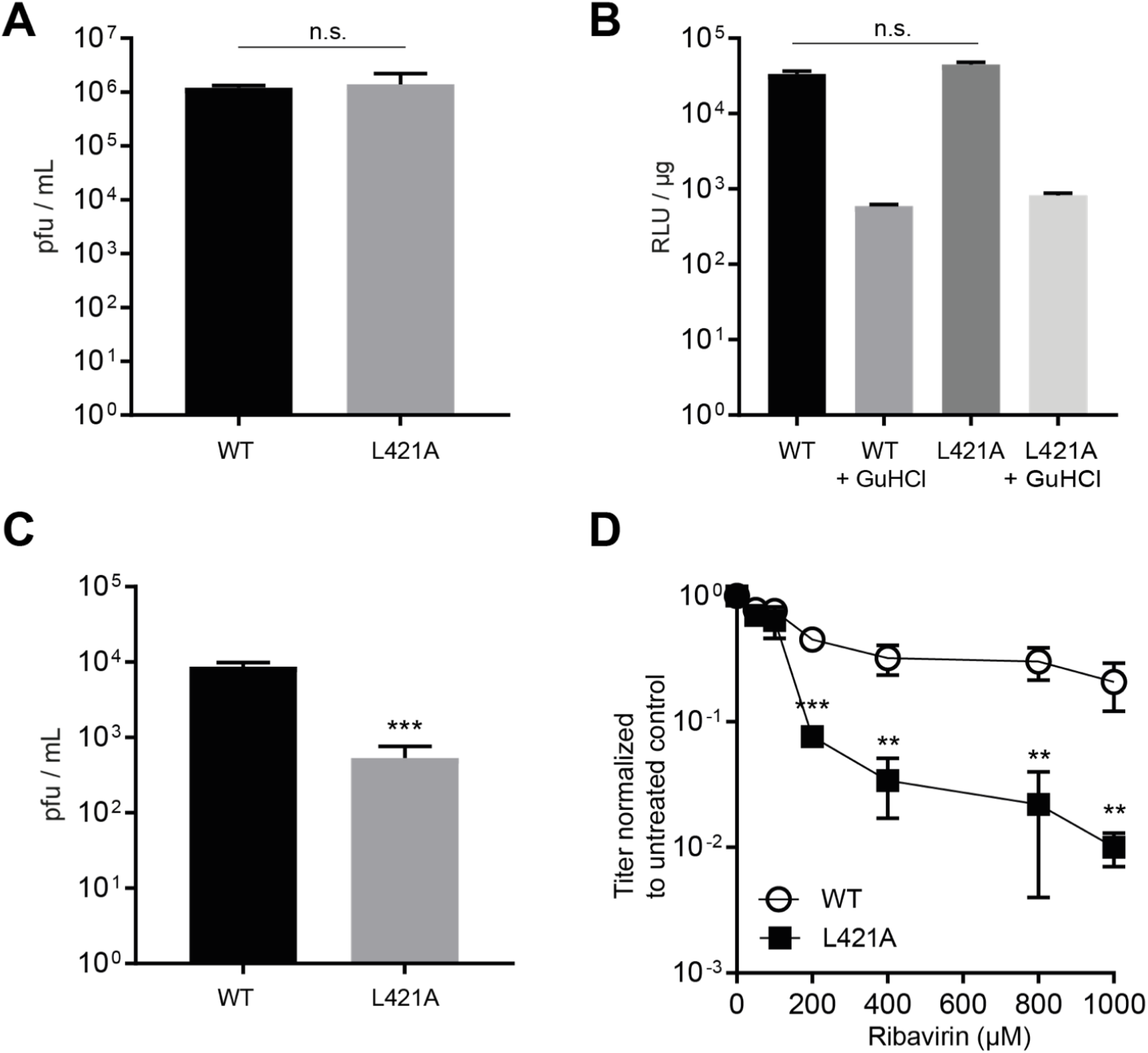
Mutation to the donor RdRp inhibits recombination and increases susceptibility to Ribavirin. **(A**) L421A mutation does not impact virus yield. Yields of virus are shown for wild-type EV-A71 C2-MP4 and the L421A variant following transfection of RNA (pfu/mL ±SD; *n = 3)*. **(B)** L421A mutation does not impact donor template replication. Cells were transfected with 250 ng of wild type EV-A71 replicon and the L421A variant ± 4mM guanidine hydrochloride. Luciferase activity is reported in relative light units *(RLU)* per microgram of total protein in the extract 8 h post transfection. **(C)** L421A inhibits EV-A71 replicative recombination. Yield of recombinant virus following transfection of either wildtype or L421A variant donor template with acceptor RNA in RD cells (pfu/mL ±SD; *n = 3)*. Statistical analyses were performed using unpaired, two-tailed t-test (***significance level p = 0.0003). **(D)** EV-A71 L421A population is highly susceptible to Ribavirin. RD cells were infected at a MOI 0.1 with wildtype or L421A variant EV-A71 C2-MP4 virus in the presence of various concentrations of Ribavirin. Following CPE, virus supernatant was clarified and used for plaque assay. Results show titer of virus normalized to an untreated control (pfu/mL ±SD; *n = 3)*. Statistical analyses were performed using unpaired, two-tailed t-test (***significance level p = 0.0004, ** p = < 0.005).

### Use of alternate donor templates significantly impacts recombinant virus yield in a predictive manner

The main goal of developing the EV-A71 recombination assay was to test its predictive power in reproducing what is observed in nature. In order to do this, two additional sub-genomic replicons (donor templates) were engineered and introduced into the assay. Importantly, both donor templates were developed from the current circulating clinical isolates TW-00073-2012 and TW-50144-2013, with phylogenetic analysis placing them within the C4 and B5 subgenogroups respectively (Fig. 3). At the RNA level, the C4 and B5 replicons share 80.3% and 79.3% sequence homology with the C2-MP4 acceptor template respectively within the region of recombination (Table 1). Although, the majority of nucleotide differences are at the wobble-base position as amino acid homology to the acceptor template is ~95% for both replicons (Table 1). Crucially, current phylogenetic data shows the C4 group has evolved by genetic shift through intra- and intertypic recombination events. Indeed, the C4 subgenotype is characterized by a higher similarity to the prototype CV-A16 virus (G-10) at the P2 and P3 region. (Fig. 3, marked in green). In contrast, analysis of B5 members clusters them in an independent clade within the genotype B group and suggests that evolution has been limited to genetic drift only (Fig. 3, arrow). As the B5 subgenogroup is not associated with current circulating recombinant viruses, we hypothesized that the B5 donor template in our recombination assay would produce significantly lower recombinant yield when compared to the C2 and C4 donor templates. A single-step growth curve for the EV-A71 C4 and B5 clinical isolates indicated no significant difference in replication (Fig. 4A). In addition, we tested the replication kinetics of the new sub-genomic replicons using a luciferase time-course assay (Fig. 4B). The results show similar luciferase signal as a function of time for all three donor templates, with the B5 replicon producing a marginally higher reporter signal at each time-point when compared to the C2 and C4 strains. As there was no significant difference in replication between the three donor templates a subsequent recombination assay was carried out. Quantification of recombinant virus showed that alternate donor templates did indeed significantly impact viable recombinant yield (Fig. 4C). All combinations tested produced detectable virus (C2/C2 = 6.9 x 10^3^ pfu/mL ± 2.4 x 10^3^, C4/C2 = 4.2 x 10^2^ pfu/mL ± 10^2^, and B5/C2 = 5.6 x 10^1^ pfu/mL ± 10) and all viruses produced were recombinant (see representative sequences in Fig. 4DE). If RNA sequence homology is the driving force for RdRp mediated ‘ copy-choice’ recombination then the significantly higher yield for the C2/C2 partners compared to the other two conditions is not surprising, given the sequence homology between donor and acceptor templates is >99%. However, the C4 and B5 replicons shared similar divergence in RNA sequence to the acceptor, but the yield of recombinant virus was significantly different. This result indeed followed what is observed in nature and supported our working hypothesis. We concluded that subtle underlying differences at either the RNA, or proteome level inhibit recombination with B5 subgenogroup members. On-going experimentation is underway to characterize these differences.

**Figure 3.**
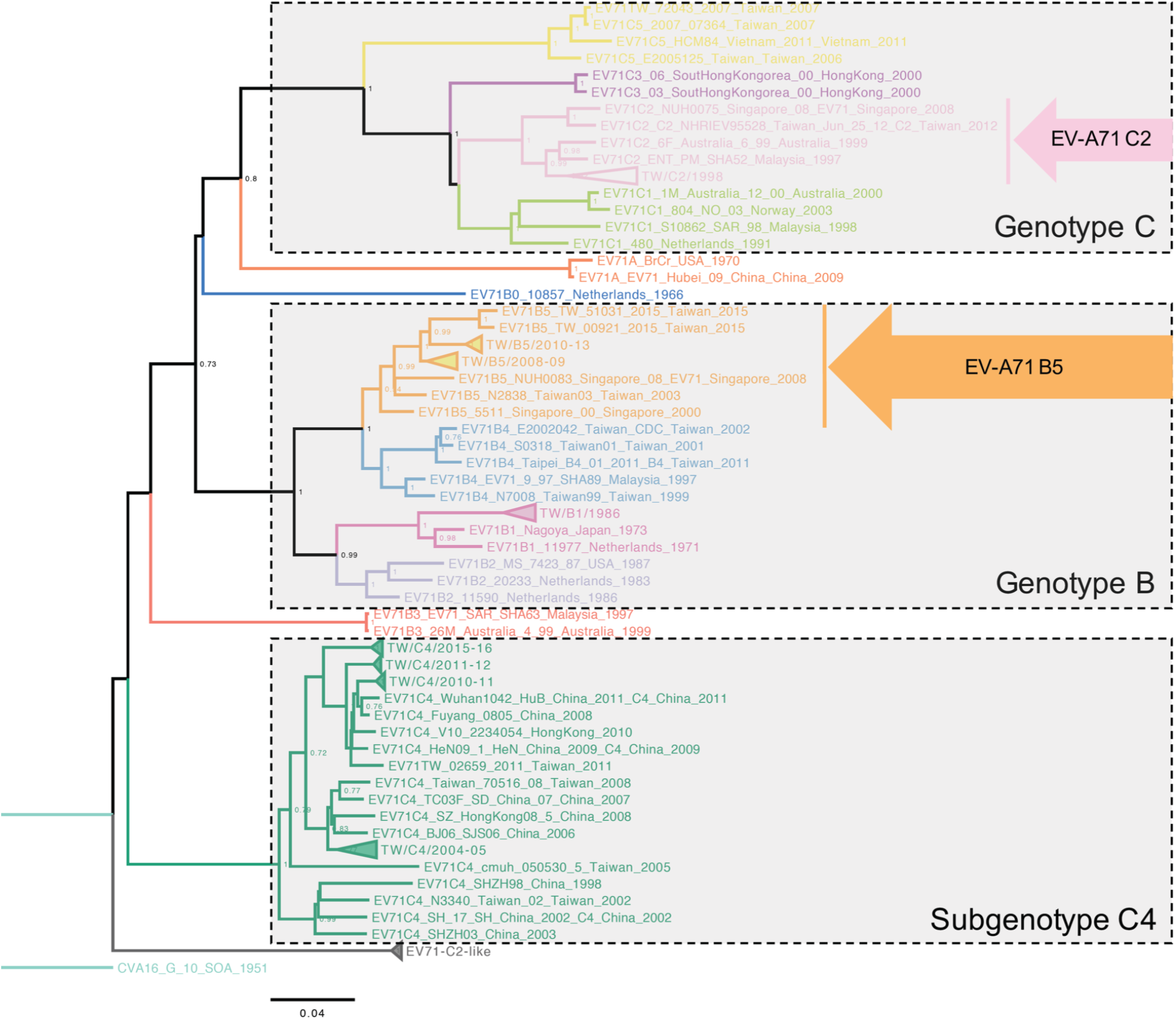
Phylogenetic analysis of EV-A71 genotypes B and C. The Neighbor-Joining phylogenetic analysis of EV-A71 genotypes B and C was based on their P2-P3 genome region, rooted by the coxsackievirus (CV) A16 prototype strain G10 (isolated in 1951). The subtrees show mixed clusters of evolutionary intra- and intertypic recombination events of analyzed EV-A71 sequences (n = 182). EV-A71 subgenotypes and genotypes depicted in different colors. The subgenotype B5 (marked by a yellow arrow) locates within the genotype B cluster, showing a ladder-like evolutionary scale. In contrast to other subgenotypes of genotype C, the subgenotype C4 (labeled in green) forms an outgroup of genotype C, closing to other recombinogenic EV-A71 strains (e.g., B3 and C2-like) and the prototype CV-A16 sequence. The probability of replicate trees in which the associated taxa clustered together in the bootstrapped data (1,000 replicates) are shown next to the branches. The phylogenetic tree is drawn to scale, with branch lengths representing the number of base substitutions per site.

**Figure 4.**
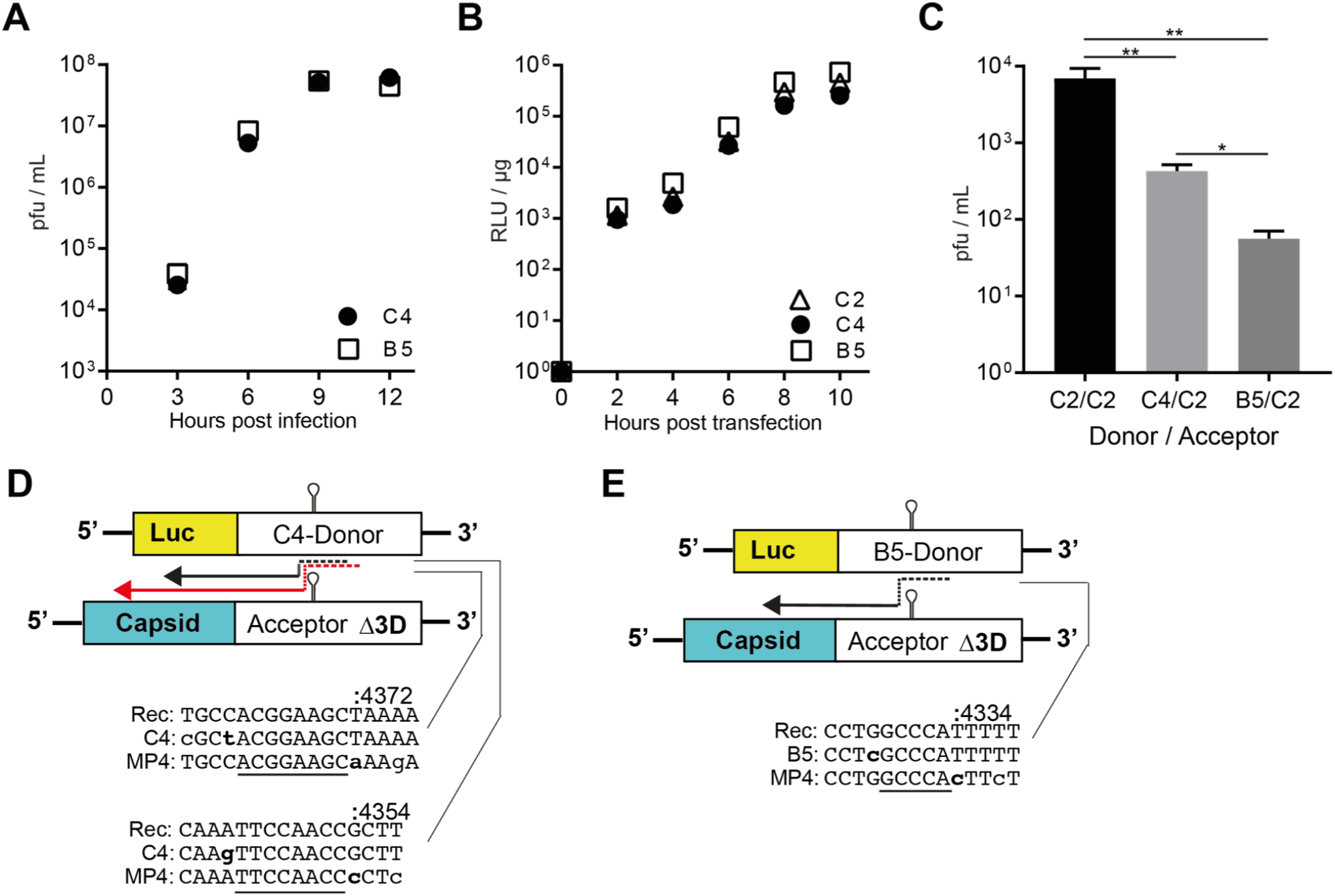
Alternate donor templates significantly impact viable recombinant frequency in a predictable manner. (A) Single-step growth curve at a MOI 10 of EV-A71 C4 and B5 strains shows no significant difference in replication.(B) C2, C4 and B5 sub-genomic replicon firefly luciferase time-course. Cells were transfected with 250 ng of each respective EV-A71 replicon and luciferase activity is reported in relative light units *(RLU)* per microgram of total protein in the extract. **(C)** EV-A71 recombination assay. RD cells were transfected with the various EV-A71 sub-genomic replicon donor and EV71Δ3D RNA. Results show yield of recombinant virus (pfu/mL ±SD; *n = 3)*. Statistical analyses were performed using unpaired, two-tailed t-test (** significance level p = < 0.01, * = < 0.05). **(D + E)** Representative recombinant sequences of plaque-purified recombinant virus from C4/C2 (E) and B5/C2 (F). Dashed arrows indicate predicted paths of viral RdRp upon template switching. Numbering refers to position upon the acceptor templates. Lower case, bold nucleotides indicate the 5’ and 3’ boundaries of recombination. Underlined sequences indicate region of homology.

**TABLE 1.** Nucleotide and amino acid identity between donor template (C4 and B5 strains) and acceptor template (C2 strain). Consensus sequences of the C4 and B5 subgenotypes were used to perform the pairwise sequence alignments with the C2-MP4 acceptor template strain. Sequence identities at the RNA and amino acid level are denoted as percentage.

|  |  | Identity (%) |  |  |  |  |  |  |  |  |  |  |  |
| --- | --- | --- | --- | --- | --- | --- | --- | --- | --- | --- | --- | --- | --- |
| EV-A71 strain | Sequence type | Overall | VP4 | VP2 | VP3 | VP1 | 2A | 2B | 2C | 3A | 3B | 3C | 3D |
| C4 | Nucleotide | 83.3 | 88.9 | 91.3 | 91.9 | 90.5 | 84.9 | 74.7 | 79.0 | 76.4 | 75.8 | 75.8 | 77.9 |
|  | Amino acid | 96.7 | 100.0 | 100.0 | 99.6 | 98.7 | 97.3 | 92.9 | 96.7 | 94.2 | 90.9 | 94.0 | 93.9 |
| B5 | Nucleotide | 80.1 | 80.7 | 81.2 | 82.0 | 83.1 | 81.2 | 75.8 | 79.7 | 78.1 | 80.3 | 77.3 | 79.1 |
|  | Amino acid | 95.5 | 100.0 | 97.6 | 97.1 | 97.3 | 96.7 | 91.9 | 95.7 | 93.0 | 95.5 | 91.8 | 93.7 |

### Isolated recombinant viruses show homology at the recombination junction(s). The trigger for recombination is sequence independent

The exact location of recombination from the primary C2/C2 recombinant viruses isolated in this study was impossible to identify due to the high sequence similarity of the two parental templates (Fig. 1C). As the C4 and B5 templates had ~20% divergence at the RNA level within the P2 and P3 regions, where template-switching would occur (Table 1), identification of recombinant junction would in principle be easier. Of note, only one independent B5/C2 recombinant sequence was identified (Fig. 4D). As the yield of recombinant virus from this combination was very low (mean of 5.6 x 10^1^ pfu/mL) this may be unsurprising and may represent the sole recombinant in the population. The C4/C2 intertypic pairing provided more viable recombinants which were subsequently used for sequence analysis. Plaque purified viruses were characterized by RT-PCR analysis in a window between the end of P1 and P2 (Fig. 5A). All recombinant viruses identified were derived from the parental genomes, as expected. In all examples, a region of between 5-11 nucleotides of shared homology between donor and acceptor templates was identified at the recombination junction with no insertions or deletions (Fig. 5A). The identified homologous sequences shared a sequence homology of only 58 ±2% and were subject to de novo sequence motif search to identify possible conserved sequences that may trigger the observed recombination events. Potentially, the cell-based recombination assay only provides information of secondary recombination products that don’ t necessarily represent the primary recombinant product, we hypothesized that those may still share similar sequence motifs, if any exist. We first analyzed the identified sequences with M-COFFEE *(38)*, a multiple sequence alignment algorithm that allows gaps between sequence regions (Fig. 5B) to allow best alignment. The computation was set to combine the following alignment algorithms: MAFFT, ClustalW, DIALIGN-TX, POA, MUSCLE, T-COFFEE, PCMA, and PROBCONS. The algorithm was not able to identify a highly conserved sequence motif and the proposed consensus sequence with a probability of 70% only exhibited few guanosines as conserved nucleotides within the sequences. Further analysis with MEME (Fig. 5C) *(39)*, which searches for ungapped sequence motifs, also failed to identify a sequence-dependent recombination trigger (E-value = 0.06). The position-dependent nucleotide probabilities (Fig. 5D) exhibit, similar to the M-COFFEE results, that guanosines are most abundant in the homologue sequences. This result gave rise to the assumption that G:C-rich sequence regions or CpG/ApU dinucleotide bias may trigger RdRp template-switching, as previous studies suggest *(40, 41)*. To evaluate this hypothesis, we compared the homologous and donor template sequences in regard to A:U and G:C nucleotide density (Fig. 5E), the number of successive G:C and A:U basepairs (Fig. 5F), and CpG density (Fig. 5G). The analyses showed no significant differences between the donor template nucleotide composition and the sequences identified at the recombination junctions. Taken together, the sequence analysis results yielded no indication of any sequence-dependent recombination trigger.

**Figure 5.**
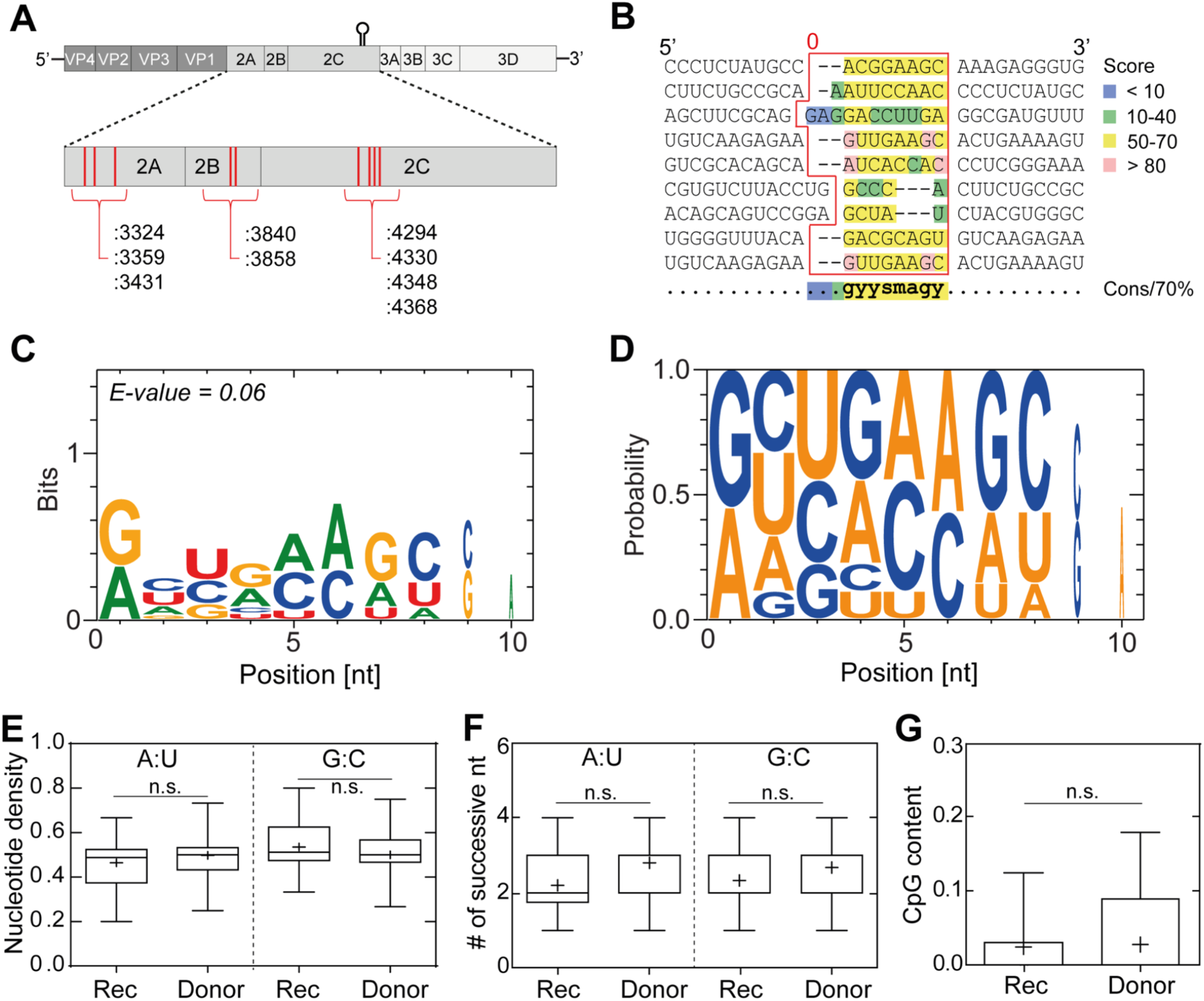
Intertypic recombination between EV-A71 C2 and C4 subgenotypes requires homology at the recombination junction but the triggers for template-switching are sequence-independent. (A) Positions of template-switching events observed during intertypic recombination between the EV-A71 C2 and C4 subgenotypes. The individual positions of observed strand switching across the P2 genome region are marked in red, including their corresponding nucleotide sequence number in respect of the C2 acceptor strand. **(B)** Sequences of bona fide C2/C4 recombinant viruses. The red border highlights the matching homologous sequences at the recombination sites with lengths between 5-11 nucleotides (x□ = *7 ± 2 nt)*. The sequences were subject to M-COFFEE, a multiple sequence alignment algorithm to identify possible gapped sequence motifs. Scores < 50 are considered to exhibit poor sequence consistency (Score 0-100). Sequence homology for the matching homologous sequences was found to be 58 ±2 %. A consensus sequence with 70% probability is shown below using IUPAC nomenclature. **(C)** Logo of ungapped de novo sequence motif search using the MEME algorithm that represents a sequence-aligned, position-dependent nucleotide-probability matrix. The resulting motif sequence with an E-value of 0.06 and bit value <1 show no statistical significance and thus failed to find a sequence motif as a recombination trigger. **(D)** Probability of position-dependent nucleotides at the homologous recombination sequences without sequence alignment. **(E)** G:C nucleotide density, **(F)** number of successive G:C/A:U bases, and **(G)** CpG content of the homologous recombination sequences and the entire C2 genome (10 nt sequence window) exhibit no significant difference. Statistical analysis was performed using oneway, two-tailed analysis of variance (ANOVA) with comparative Tukey post-hoc test *(n.s.= non-significant)*.

## Discussion

Though EV-A71 infection generally causes mild diseases like HFMD in children, it can lead to severe cardiorespiratory and neurological complications *(42)*. Intratypic recombination between circulating strains of EV-A71 and intertypic recombination with species A Coxsackievirus produces the outbreak-associated strains of EV-A71, which exhibit increased virulence and/or transmissibility *(1, 43)*. The subsequent human mortality and morbidity associated with such events can be substantial *(44)*. All of our current understanding of recombination in the generation of new strains of enteroviruses are based upon retrospective phylogenetic analysis which shows us that recent intra- and intertypic EV-A71 recombination events are limited to members of the same species group *(14-16, 42)*. In addition, the production of chimeric enterovirus genomes in other studies indicates a high level of plasticity *(45-47)*. This plasticity, however, seems to be limited to intraspecies members, as no recent evidence for interspecies enterovirus recombination has been documented. Is this just due to co-circulation? Or, is it due to genomic / proteomic compatibilities that are only available with other group members?

Ongoing studies using the prototypical species C enterovirus, PV, have provided unique insights into the potential triggers and mechanisms of recombination *(20, 21)*. In general, all recent publications support the notion of replicative RdRp-mediated recombination as the primary source of new virus hybrid genome *(10, 19-21)*. Our major aim was to use this knowledge to develop an assay that would allow prediction of recombination between current circulating EV-A71 strains. This study reports the development of the first non-polio enterovirus recombination assay that will allow for the continued study of recombination in this medically important group of viruses. As recombination frequencies in the closely related PV have been shown to occur at between 10^-4^ - 10^-5^ *(6, 48)* any impact on overall replication of the virus would impact any recombination event. RD cells produced the highest luciferase signal from the C2-replicon donor template (a surrogate for replication) following transfection when compared to Vero cells (Fig. S2) and were therefore chosen for all subsequent experiments. The cell-based recombination assay shows that recombination in EV-A71 is primarily replicative (Fig. 1), as only minimal viable recombinant virus was identified from the non-replicative partners. We therefore propose that the recombination process is RdRp mediated and mechanistically similar to PV *(10, 20, 21)* and therefore different to the previously reported non-replicative mechanisms observed in PV by Gmyl and co-workers (32, *33)* and other RNA viruses like Hepatitis C *(34)*. In support of this interpretation are the results shown in figure 2. The L420 residue in the RdRp of PV is in a region of the polymerase that directly interacts with the viral RNA (21). This residue is conserved in EV-A71 and is located at position L421. Studies in PV have shown that a L420A mutation can inhibit replicative recombination by ~100-fold, while having no impact upon replication *(21)*. The same mutation in EV-A71 also produced a similar phenotype. Recombination was significantly reduced while having no impact upon replication of the full-length virus or replicon donor template (Fig. 2). The structure of the RdRp from PV and EV-A71 are very similar and many key residues are conserved. The observation of reduced recombination from the same mutation to a similar region upon the RdRp is suggestive of conservation in mechanism. In support of this a similar mutation of the Gly-64 residue that has been shown to be important for fidelity in PV have also been engineered into EV-A71 with a similar outcome *(49, 50)*. Current opinion proposes that enteroviruses have evolved to recombine in order to overcome the deleterious impacts of high mutation rates in order to maintain population fitness *(36, 37)*. Or, alternatively it may be as a consequence/biproduct of replication speed *(51)*. In either circumstance a reduction in recombination rate should negatively impact the fitness of the viral population. The L421A mutation led to the EV-A71 population being significantly more susceptible to the nucleoside analogue mutagen Ribavirin than wildtype (Fig. 2D). Again, this phenotype is conserved in PV.

We introduced additional circulating EV-A71 partners (subspecies C4 and B5) (Fig. 4). Both shared similar RNA sequence similarity to the acceptor template in the P2 and P3 regions at ~80% and ~95% at the amino acid level (Table 1). However, the phylogenetic tree (Fig. 3) of the C4 strain shows it has evolved by genetic drift and shift. In contrast, the B5 phylogeny (Fig. 3) currently shows no evidence of shift, with evolution being limited to genetic drift only. The yield of C4/C2 recombinant was significantly lower than the C2/C2 pairing. This was not surprising as similar observations have been seen with the intertypic PV1/PV3 CRE-REP partners *(19)*, and are presumably a reflection of the RNA sequence divergence. What cannot be explained by sequence divergence is the significantly lower B5/C2 recombinant yield. Is this due to lack of opportunity? i.e. distinct sites of RNA replication within the cell which decrease the likelihood of mixed replication complexes where RdRp mediated template-switching can occur *(52)*. Or, as a result of a non-functional proteome following recombination? Potentially, recombinant RNA is being formed but may be non-infectious following packaging i.e. genome may be unable to replicate due to incorrect polyprotein processing, or may lack suitable protein-protein interactions required for packaging *(53)*. This hypothesis is supported by phylogenetic analysis of isolated circulating EV-A71 recombinant virus, which suggest functionality of the encoded polyprotein is the key determinant of viability, as recombination ‘ hot-spots’ primarily localize to gene boundaries within the non-structural region *(12, 42, 54)*. Experimentation is currently underway to identify the limitations to B5/C2 recombination. However, and most importantly, these observations represent what is currently being observed in nature; the B5 strain is circulating as a pure lineage and is not a recombinant strain.

The recombinant viruses that were isolated from the C4/C2 and B5/C2 recombination assays are suggestive of a ‘ copy-choice’ mechanism of recombination where sequence homology between the two parental templates drives the template-switch event *(55)*. All had regions of homology that were between 5-11 nucleotides in length (Fig. 5). The triggers of the template-switch itself seem to be sequence independent. Historical studies of circulating PV recombinant virus identified an ApU dinucleotide bias immediately prior to the recombination junction *(56)*. Alternatively, cell-based studies suggested that RNA structure and GC-rich regions are triggers for RdRp template-switching *(41)*. However, these observations are not transferrable to EV-A71, as our analyses of possible conserved sequence motifs and nucleotide composition at the identified recombination junctions yielded no indication of sequence-dependent template-switching. Although, all historical analysis of circulating recombinant viruses are based upon evolved genomes that may not necessarily represent the primary product of recombination. Indeed, recent studies in PV suggest that recombination occurs in a biphasic fashion where promiscuous, sequence-independent template-switching occurs that is then followed by secondary selection of the most ‘ replication competent’ viruses *(19)*. Conceivably, the sequences we have identified from our C4/C2 intertypic pairing are the primary product of recombination that have yet to evolve by natural selection.

We believe that our described EV-A71 specific recombination assay and the results within will provide the basis for the further dissection of this key evolutionary process. Further, we propose that this cell-based recombination assay has the ability to predict recombination between the current circulating EV-A71 strains that are of public health relevance.

## Materials and Methods

### Cell culture

Adherent monolayers of African green monkey (Vero) and human embryonic rhabdomyosarcoma (RD) were grown in Dulbecco’ s Modified Eagle Medium (DMEM). Media was supplemented with 100 U/mL penicillin, 100 μg/mL streptomycin, and 10% Heat Inactivated (HI)-FBS. All cells were passaged in the presence of trypsin-EDTA. Where stated, guanidine hydrochloride (Sigma) was added to growth media at 4mM. Wild-type and recombinant EV-A71 virus was recovered following transfection of RNA generated *in vitro* (see below) from full-length cDNA, or from recombination assay parental partners. Virus was quantified by pfu/mL.

### Plasmids, in vitro transcription, cell transfection and recombinant virus quantification

The mouse adapted EV-A71 C2-MP4 infectious clone *(30)* was kindly provided by Dr. Jen-Reng Wang (Cheng Kung University, Taiwan) and modified by insertion of a ribozyme sequence between the T7 promoter and viral genome sequence in a pBR-derived plasmid. The EV-A71 C2 replicon was modified from a previously described EV-A71 C2-2231 replicon *(22)* by addition of a T7-ribozyme and polyA sequence inserted at the 5’ and 3’ end of the replicon sequence in a pBR-derived plasmid. The EV-A71 C4 replicon was constructed from the infectious clone, which was derived from the clinical strain TW-00073-2012. Full-length genome with 5’ T7/ribozyme and 3’ polyA were amplified *via* PCR and cloned into the pCRII-TOPO vector (Thermo Fisher Sci.). In order to construct the replicon, a cassette consisted of the SacII restriction enzyme site and 2Apro cleavage site was inserted into the 5’ UTR/VP1 and VP4/2A boundary, respectively. The P1 fragment of the cassette-containing plasmid was later replaced by the luciferase coding sequence by SacII restriction enzyme digestion. A similar strategy was utilized to construct the B5 infectious clone and sub-genomic replicon based on the sequence of clinical strain TW-50144-2013 *(54)*. The EV-A71A3D template was constructed from the full-length EV-A71 C2-MP4 infectious clone by removal ~ 800 nt between the blunt cutting restriction sites (ScaI and *NruI)* within the 3D^po1^ coding region. The C2-ΔIRES-replicon was constructed by removal of the majority of the IRES region between two ApaI restriction sites located at positions 37 and 767 of the EV-A71 C2 replicon template. The C2-L421A mutant replicon and infectious clone were constructed by using site-directed mutagenesis. The EV-A71 C2 replicon and C2-ΔIRES-replicon were linearized with *SalI*. The EV-A71-MP4, EV-A71Δ3D cDNA were linearized with *EagI*. The EV-A71 C4 and B5 replicon was linearized with *NotI*. All linearized cDNA was transcribed *in vitro* using T7 RNA Polymerase treated with 2U DNAse Turbo (ThermoFisher) to remove residual DNA template. The RNA transcripts were purified using RNeasy Mini Kit (Qiagen) before spectrophotometric quantification. Purified RNA in RNase-free H_2_O were transfected into cell lines using TransMessenger (Qiagen). The mixture was incubated according to the manufacturer’s instructions and added to RD cell monolayers in 12-well tissue culture plates. Virus amount was quantified by plaque assay. Briefly, media supernatant and cells were harvested at time-points post transfection (specified in main text), subjected to three freeze-thaw cycles and clarified. Supernatant was then used on fresh RD cells in 12-well plates, virus infection was allowed to continue for 30 min. Media was then removed, and cells were subjected to 2x PBS (pH 7.4) washes before a 1% (w/v) agarose-media overlay was added. Cells were incubated for 3-4 days and then fixed and stained with crystal violet for virus quantification.

### Ribavirin sensitivity assay

RD cells were treated with ribavirin for 3 hrs before infection (doses specified elsewhere). Ribavirin treated cells were then infected at MOI 0.1 with either wild-type of L421A variant of EV-A71-C2-MP4. Following infection cells were washed with PBS x3 and media was replaced with ribavirin. Infection was allowed until CPE was observed. Cells and supernatant were freeze/thawed x 3 and media was clarified and used for plaque ass

### Single-step growth curve of the EV-A71 C4 and B5 full-length viruses

RD cells in 6-well plates were infected by each virus at a MOI of 10 in serum-free media. One hour later, cells were extensively washed by PBS and refreshed in 2% serum-containing media. Virus was harvested at different time-points post infection and the virus yield was quantified by plaque assay.

### Luciferase assays

Supernatant was removed from transfected cell monolayers, and cells were briefly washed with PBS and lysed using 100 μl 1x Glo Lysis Buffer (Promega®) per well in a 12-well plate. The oxidation reaction was catalyzed by the addition of 10μl cell lysate to 10μl room temperature *Bright-Glo* Luciferase Assay System (Promega®) substrate. Luciferase activity was measured using a luminometer with values normalized to protein content of the extract using a protocol as described previously *(57)*.

### Recombinant virus sequencing

Recombinant viruses were isolated from individual plaques by incubating the media/agar plug overnight in 1x PBS. Viral RNA was isolated using Qiagen viral RNA isolation kit, following the manufacturers protocol. RNA was reverse transcribed with an oligo-T primer using Superscript II (Invitrogen) following the manufacturers protocol. PCR was carried from the P1 region of the acceptor template to the end of the P2 region of the donor using Phusion high-fidelity DNA polymerase (NEB) following the manufacturers protocol. PCR products were gel purified, A-tailed and sub-cloned into a pCRII-TOPO vector (Thermo Fisher Sci.) for sequencing. Clustal Omega was used for sequence alignment to identify recombinant junctions.

### Phylogenetic analysis of the P2-P3 region of the EV-A71

The evolutionary history of EV-A71 and its relationship with CV-A16 was inferred using the Neighbor-Joining method and constructed in MEGA7 *(58)*. Sequences including 182 EV-A71 sequences and the prototype sequence of CV-A16 were analyzed and rooted with the CV-A16 prototype strain G10 isolated in 1951. To specifically discriminate the recombinogenic property of different genotype/subgenotype of EV-A71, P2-P3 rather than the VP1 region was analyzed. The probability of replicate trees in which the associated taxa clustered together was determined from bootstrapped data (1,000 replicates) *(59)*. The evolutionary distances were computed via MEGA7 using the Jukes-Cantor method *(60)* and are in the units of the number of base substitutions per site. All positions with less than 95% site coverage were eliminated.

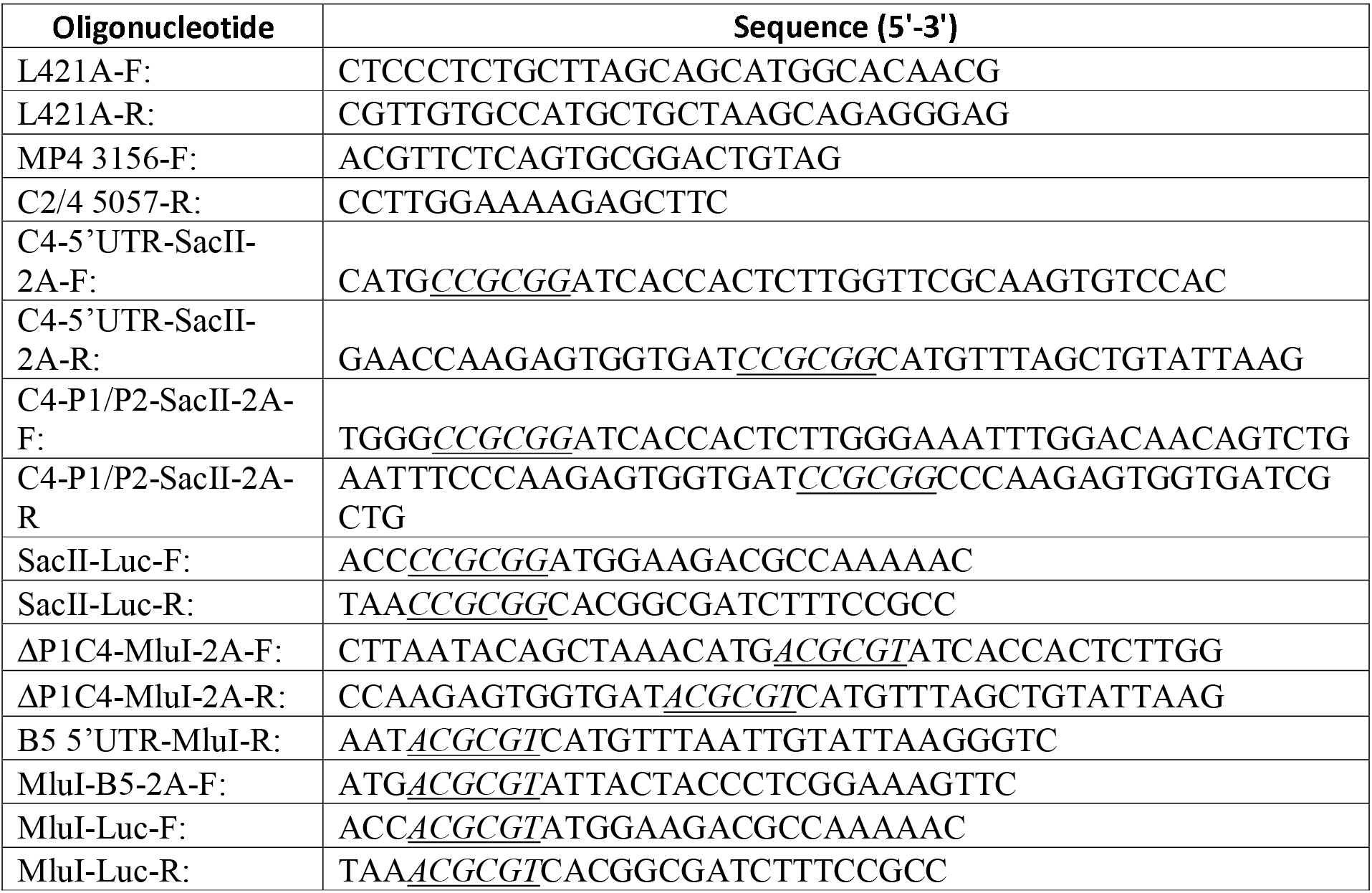
Oligonucleotides used in the study.

## Acknowledgements

Funding to CEC, ND and SRS via grant (RGP0011/2015) from the Human Frontier Science Program (HFSP). Work was further funded by NIH grant no. R01AI45818 to CEC. In addition, SRS was also supported by grants from the Ministry of Science and Technology (MOST), Taiwan (MOST 107-3017-F-182-001), and the Research Center for Emerging Viral Infections from The Featured Areas Research Center Program within the framework of the Higher Education Sprout Project by the Ministry of Education (MOE) in Taiwan.

---

**Supplemental figure 1: Engineering a hammerhead ribozyme into the EV-A71 C2-replicon.**

A hammerhead ribozyme was engineered immediately 5’ of the EV-A71 internal ribosomal entry sequence (IRES). This ensures an authentic 5’ sequence following T7 RNA polymerase transcription. Luciferase signal was measured at 8 h following transfection into RD cells and normalized to protein content. Result shows a 3Log_10_ improvement in luciferase signal (a surrogate for replication). Statistical analyses were performed using unpaired, two-tailed t-test (***significance level p = 0.0001). Error bars represent standard deviation three replicate samples.

**Supplemental figure 2: Luciferase signal following transfection of C2-replicon into RD and Vero cells**

Luciferase signal was measured at 8 h following transfection into RD and Vero cells and normalized to protein content. Result shows a near 2Log_10_ higher luciferase signal (a surrogate for replication) in RD cells when compared to Vero. Statistical analyses were performed using unpaired, two-tailed t-test (*** significance level p = 0.0001). Error bars represent standard deviation three replicate samples.

